# Systemic deficits in lipid homeostasis promote aging-associated impairments in B cell progenitor development

**DOI:** 10.1101/2024.09.26.614999

**Authors:** Silvia Vicenzi, Fangyuan Gao, Parker Côté, Joshua D. Hartman, Lara C. Avsharian, Ashni A. Vora, R. Grant Rowe, Hojun Li, Dorota Skowronska-Krawczyk, Leslie A. Crews

## Abstract

Organismal aging has been associated with diverse metabolic and functional changes across tissues. Within the immune system, key features of physiological hematopoietic cell aging include increased fat deposition in the bone marrow, impaired hematopoietic stem and progenitor cell (HSPC) function, and a propensity towards myeloid differentiation. This shift in lineage bias can lead to pre-malignant bone marrow conditions such as clonal hematopoiesis of indeterminate potential (CHIP) or clonal cytopenias of undetermined significance (CCUS), frequently setting the stage for subsequent development of age-related cancers in myeloid or lymphoid lineages. At the systemic as well as sub-cellular level, human aging has also been associated with diverse lipid alterations, such as decreased phospholipid membrane fluidity that arises as a result of increased saturated fatty acid (FA) accumulation and a decay in n-3 polyunsaturated fatty acid (PUFA) species by the age of 80 years, however the extent to which impaired FA metabolism contributes to hematopoietic aging is less clear. Here, we performed comprehensive multi-omics analyses and uncovered a role for a key PUFA biosynthesis gene, *ELOVL2*, in mouse and human immune cell aging. Whole transcriptome RNA-sequencing studies of bone marrow from aged *Elovl2* mutant (enzyme-deficient) mice compared with age-matched controls revealed global down-regulation in lymphoid cell markers and expression of genes involved specifically in B cell development. Flow cytometric analyses of immune cell markers confirmed an aging-associated loss of B cell markers that was exacerbated in the bone marrow of *Elovl2* mutant mice and unveiled CD79B, a vital molecular regulator of lymphoid progenitor development from the pro-B to pre-B cell stage, as a putative surface biomarker of accelerated immune aging. Complementary lipidomic studies extended these findings to reveal select alterations in lipid species in aged and *Elovl2* mutant mouse bone marrow samples, suggesting significant changes in the biophysical properties of cellular membranes. Furthermore, single cell RNA-seq analysis of human HSPCs across the spectrum of human development and aging uncovered a rare subpopulation (<7%) of CD34^+^ HSPCs that expresses *ELOVL2* in healthy adult bone marrow. This HSPC subset, along with *CD79B*-expressing lymphoid-committed cells, were almost completely absent in CD34^+^ cells isolated from elderly (>60 years old) bone marrow samples. Together, these findings uncover new roles for lipid metabolism enzymes in the molecular regulation of cellular aging and immune cell function in mouse and human hematopoiesis. In addition, because systemic loss of ELOVL2 enzymatic activity resulted in down-regulation of B cell genes that are also associated with lymphoproliferative neoplasms, this study sheds light on an intriguing metabolic pathway that could be leveraged in future studies as a novel therapeutic modality to target blood cancers or other age-related conditions involving the B cell lineage.

## INTRODUCTION

The aging immune system is typified by increased bone marrow adiposity^1^ and a shift towards myeloid cell development^2^ at the expense of lymphocyte maturation.^3, 4^ This is due in part to age-related defects in hematopoietic stem cell (HSC) function^5^ that promote altered survival, dormancy and regenerative capacity of specific hematopoietic lineages, as well as age-related alterations in the bone marrow microenvironment.^6^ At the systemic as well as sub-cellular level, human aging has been associated with diverse lipid alterations, such as decreased phospholipid membrane fluidity that arises as a result of increased saturated fatty acid (FA) accumulation and a decay in n-3 polyunsaturated fatty acid (PUFA) species by the age of 80 years.^7–9^ While many studies have examined lipid membranes and the role of FA biosynthesis and metabolism in the aging or diseased brain,^9–11^ the extent to which impaired FA metabolism contributes to immune system aging and impaired hematopoietic stem and progenitor cell (HSPC) development is less clear. Notably, age-related immune dysfunction or immunosenescence has the potential to set the stage for the development of diverse pre-malignant disorders such as monoclonal gammopathy of undetermined significance (MGUS) or clonal hematopoiesis (CH) of indeterminate potential (CHIP).^12^ Therefore, studies aimed at interrogating the precise FA metabolism pathways that impair immune system function during aging could provide new insights and novel molecular targets for the prevention and treatment of clinical conditions related to aging.

Previously, we identified key molecular signatures of human HSPC aging, which included alterations in inflammation-responsive pathways and RNA splicing regulation, along with oxidative phosphorylation and metabolism-associated genes in purified aged versus young human HSPCs.^13, 14^ In addition, the development of robust models to systematically interrogate FA synthesis and lipid metabolism in organismal aging have enabled elucidation of the contribution of the long chain (LC)-PUFA elongation enzyme, Elongation of very long chain fatty acids protein 2 (*Elovl2*) to the acquisition of tissue-specific features of accelerated aging.^7, 15^ These novel models of aging biology allowed the discovery of *Elovl2* DNA methylation-dependent loss of function as a key driver of age-related defects in eye and neural tissues, and development of a unique mouse model that expresses a mutant inactive form of *Elovl2* also recapitulates these features of physiological aging, which are even more severe in mutant versus age-matched wild-type controls.^11, 16^

In mammals, *Elovl2* is exclusively responsible for the elongation of 22-carbon PUFAs to form 24-carbon PUFAs, and to a lesser extent for the elongation of 20-carbon PUFAs to 22-carbon species. While some FAs, including the omega-3 (n-3) FAs EPA and DHA can be obtained from dietary sources, LC-PUFAs containing 24 or more carbons are primarily generated *in vivo* through enzymatic elongation. 24-carbon PUFAs are present as natural products in some marine organisms, however they make up only a very minor fraction of FA supplements derived from dietary sources. Thus, alterations in the endogenous catabolism of PUFAs during systemic aging could have broad impacts on the lipid composition of diverse tissues and of cellular membranes within those tissues, as has been shown previously in the retina and in neurons.^11, 16^ Moreover, whole blood DNA methylation studies have identified *ELOVL2* promoter hypermethylation as the one of the best biomarkers of human chronologic aging,^17^ however, the role of *ELOVL2* function in blood stem cell development has not been established.

Here, we performed comprehensive multi-omics analyses along with functional studies to uncover the role of *ELOVL2* in mouse and human immune cell aging. Whole transcriptome RNA-sequencing studies of cells isolated from mouse bone marrow revealed global down-regulation in lymphoid cell markers and expression of genes involved specifically in lymphoid development occurring in *Elovl2* mutant animals compared with age-matched controls. Flow cytometric analyses of immune cell markers confirmed that an age-related loss of mature B cell markers was exacerbated in the bone marrow of *Elovl2* mutant mice, while lipidomics studies extended these findings to reveal select alterations in lipid species in aged and *Elovl2* mutant mouse bone marrow samples, suggesting significant changes in the biophysical properties of cellular membranes.

To further explore the clinical relevance of these findings and elucidate connections between lipid homeostasis and lymphoid specification in human samples, we analyzed public datasets along with a unique human cell dataset^18, 19^ to characterize *ELOVL2* expression and its correlation with lymphoid cell markers at single-cell resolution in human HSPCs spanning gestation, maturation, and aging. These analyses revealed the presence of a subset of CD34^+^ HSPCs that express *ELOVL2* in healthy adult bone marrow. This population, along with lymphoid-biased HSPCs, was almost undetectable in CD34^+^ cells isolated from elderly (> 60 years old) bone marrow samples. Together, these findings uncover new roles for lipid metabolism enzymes as key molecular regulators of immune aging in mouse and human hematopoiesis, with potential implications for anti-aging interventions (restoring ELOVL2 expression and activity) as well as novel anti-cancer strategies (down-modulating ELOVL2 expression or activity) for lymphoproliferative neoplasms.

## RESULTS

### Lymphoid-specific molecular and cellular deficits associated with bone marrow aging are accelerated in a mouse model of impaired lipid metabolism

A growing body of evidence has indicated that physiological mouse and human immune system aging is characterized by myeloid skewing in the bone marrow and underlying hematopoietic stem and progenitor cell dysfunction. However, no animal models of accelerated bone marrow aging are currently available that recapitulate these effects to enable research into the mechanisms contributing to these alterations. Here we have comprehensively characterized RNA (whole transcriptome sequencing), protein (flow cytometry), and lipid species (lipidomics) in the bone marrow of a unique mouse model of impaired lipid metabolism that has been previously shown to display features of accelerated aging in other tissues (eye and neural tissue).^11, 16^ These animals express a cysteine-to-tryptophan substitution (C234W) in the gene encoding for the lipid elongation enzyme, *Elovl2*,^11, 16^ which selectively inactivates enzymatic activity of ELOVL2 required to elongate 22-carbon PUFAs.^20^

In order to characterize the molecular profile of the bone marrow of these *Elovl2*^C234W^ (*Elovl2-*MUT) mice, we performed whole transcriptome sequencing of bone marrow cells isolated from wild-type (WT) young (2-3 months old), WT aged (18-22 months old), MUT aged (18-22 months old), and WT geriatric (27 months old) mice, including both male and female animals for analysis. For breeding-related reasons, young mutant mice were not readily available for analyses, so we first focused on evaluating the global differences in whole transcriptome sequencing data generated from strain-matched MUT aged versus WT aged mice. A total of 389 genes were found to be differentially expressed, with 91 genes upregulated, and 298 downregulated. Gene set enrichment analyses of KEGG pathway gene orthologs revealed a significant down-regulation of genes involved in cytokine receptor signaling along with immunodeficiency and B-cell receptor (BCR) signaling, which included myriad B/T/plasma cell maturation-specific surface receptors and lineage specification genes (Figure 1A-C; S1A). Among these, well-known lymphoid progenitor and mature B-cell marker genes were significantly down-regulated in MUT aged versus WT mice, including *Il7r, Cd19*, *Cd79a/b*, and *Cd22*, among other genes that were differentially expressed in the KEGG B-cell receptor signaling pathway (Figure 1C). Validation of the sequencing results through additional RNA-seq analysis in an expanded cohort of samples that included WT geriatric mice confirmed that the majority of these genes were significantly reduced in aged MUT versus WT mice (Figure 1D). Furthermore, similar trends were observed in additional samples analyzed by quantitative (q)RT-PCR, with the most severe B cell deficits occurring in geriatric mice (Figure 1D; S1B). Notably, the levels of B cell marker gene expression measured in MUT aged (18-22 m/o) mice were reduced to levels similar to those observed in WT geriatric mice of more advanced ages (27-28 m/o) (Figure 1D; S1B). This is in line with separate studies of the ELOVL2 enzyme that have evaluated other tissues and reported an approximate 6-month acceleration of age-related deficits,^16^ which we now observe also occurs in the bone marrow of mutant animals of the same strain.

**Figure 1:**
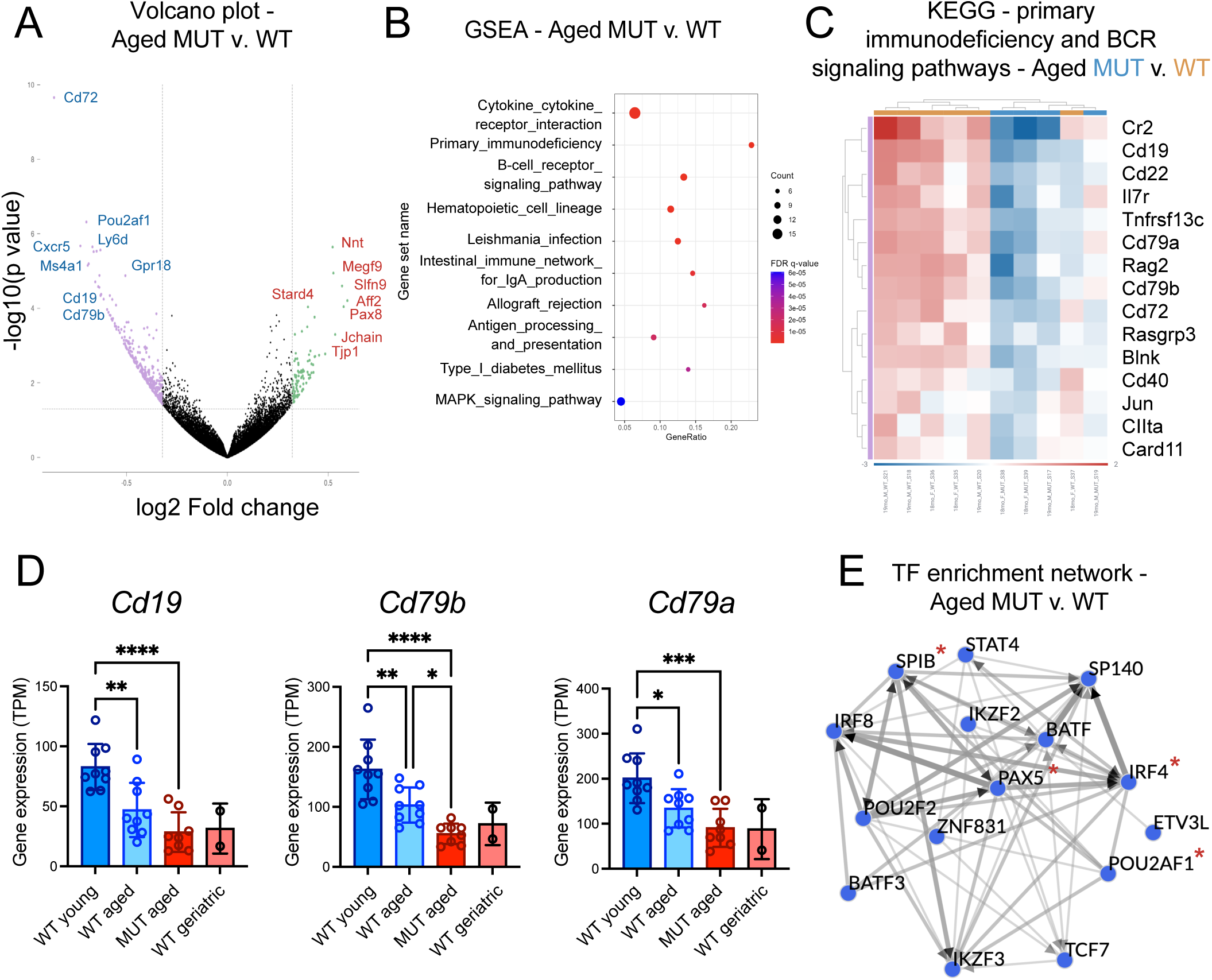
Whole transcriptome analyses delineating age-related molecular deficits in the bone marrow of a mouse model of impaired lipid metabolism. Total bone marrow cells from 18-22 month old (“aged”) WT versus ELOVL2 C234W MUT mice were analyzed by whole transcriptome RNA-sequencing with targeted genes validated in an expanded cohort of aged mice also including WT young (2-3 months old) and WT geriatric (27 months old) mice. A) Volcano plot displaying 91 upregulated and 298 downregulated genes in MUT (n=2 female, n=2 male) versus WT (n=3 female, n=3 male) aged (18-19 months old) mouse bone marrows. B) Dot plot showing the relative p values, overlapping gene counts, and gene ratios of the top 10 differentially regulated KEGG pathways by gene set enrichment analysis (GSEA) of the significantly upregulated and downregulated genes in the aged-matched MUT vs WT mouse bone marrows. C) Among the top three differentially regulated gene sets from (B), KEGG pathway gene orthologs show significant down-regulation of genes involved in primary immunodeficiency and B-cell receptor (BCR) signaling. Gene sets were combined for the heatmap display due to several overlapping genes in the two sets. D) Validation of gene expression levels from RNA-sequencing of an expanded cohort of mice including WT young (2-3 months old, (n=3 female, n=6 male)), WT aged (18-22 months old, (n=3 female, n=6 male)), MUT aged (18-22 months old, (n=4 female, n=4 male)), and WT geriatric (27 months old, (n=2 male)) samples with relative gene expression results shown as transcripts per million (TPM). For all panels, significance was determined using one-way ANOVA with parametric or non-parametric tests based on normality tests for each dataset (*p<0.05, **p<0.01, ***p<0.005, ****p<0.001). E) The ChIP-X Enrichment Analysis 3 (ChEA3)^21^ tool was applied to evaluate all of the differentially expressed genes identified in (A) and uncover potential transcription factor co-regulatory networks responsible for the gene expression changes observed in MUT aged versus WT mice.

We noted that the top genes down-regulated in the MUT v. WT aged mouse bone marrows included several transcription factors known to be vitally important to B cell and plasma cell development, including *Irf4*, *SpiB*, *Pax5*, and *Pou2af1* (also known as OCT-binding factor-1, or OBF1). We postulated that core transcriptional B cell regulatory programs might be depleted in the hematopoietic compartment of these animals. To investigate this possibility, we then performed a transcription factor enrichment analysis using a previously described computational tool, ChIP-X Enrichment Analysis 3 (ChEA3), which ranks transcription factors associated with user-provided gene sets.^21^ This analysis revealed a striking signature of interacting B and plasma cell-regulatory transcription factors that are known to function upstream of the observed differential gene expression observed in the MUT v. WT mouse bone marrows (Figure 1E). While only a few of these transcription factors were themselves differentially expressed in our dataset, many of them are known or predicted binding partners or regulators of other factors in the enrichment list, as visualized in the transcription factor enrichment network analysis (Figure 1E). For example, the transcription factors IRF4, SPIB, STAT, and BATF family members have been previously reported to interact with each other and are also regulated by other transcription factors highlighted in the network analysis (e.g., IKZF3).^22–25^ Furthermore, many of these transcription factors have been implicated in the onset or progression of age-related B and plasma cell malignancies such as multiple myeloma.^26–28^

Since our whole transcriptome analyses were performed on bulk bone marrow samples, we hypothesized that these molecular deficits might reflect a selective loss of lymphoid populations in the bone marrow of aged *Elovl2-*MUT mice compared to WT controls. In this context, multi-parameter flow cytometry analyses of B and plasma cell populations revealed a significant loss of total CD19^+^ and CD79b^+^ lymphoid lineage cells in *Elovl2-*MUT mice compared to age-matched WT controls, with a concomitant increase in CD11b^+^ myeloid cells (Figure 2A), which is known to occur in physiological aging but has not been previously linked to age-related deficits in lipid metabolism.

**Figure 2:**
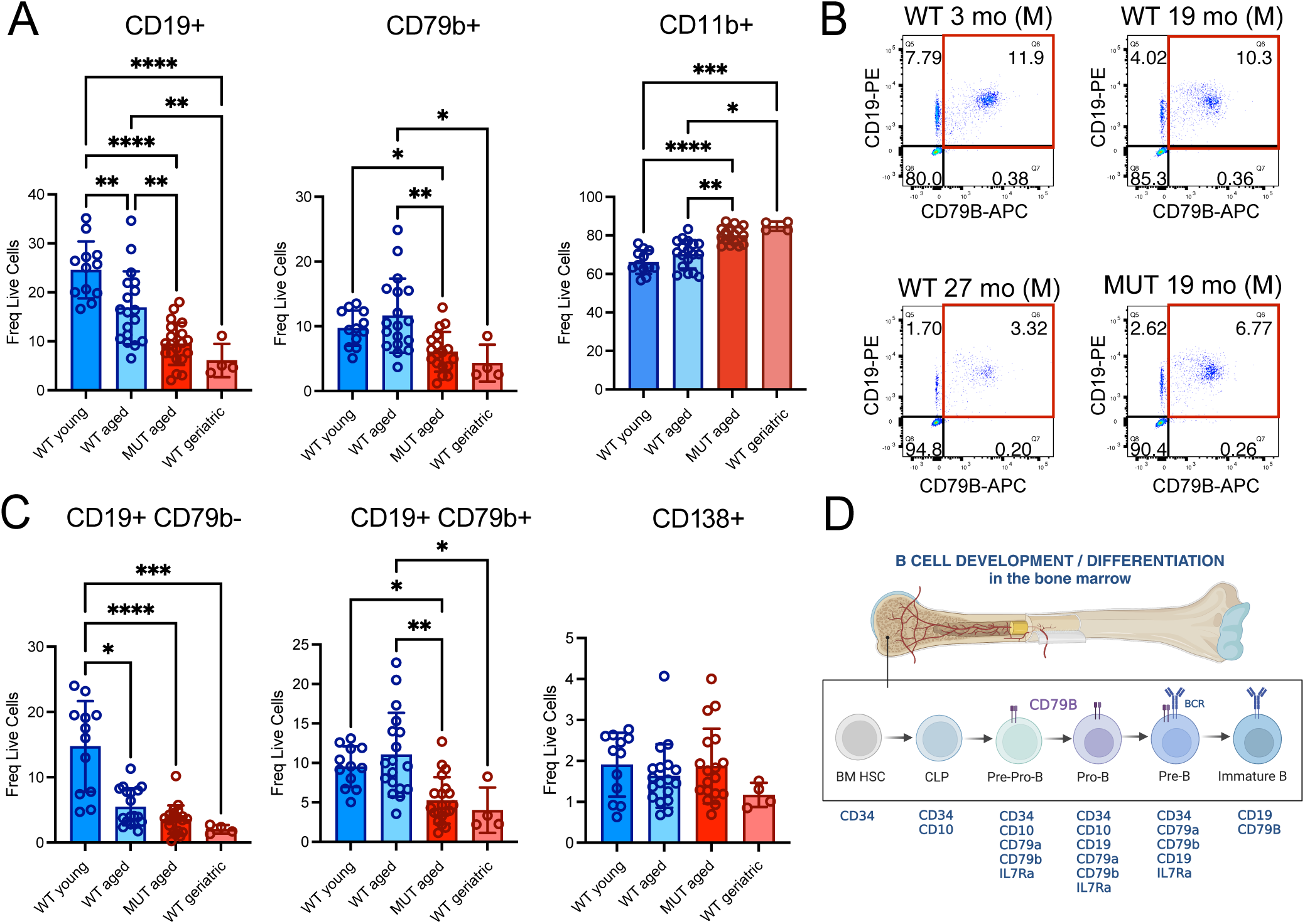
Immunophenotypic profiling of markers of B and myeloid cell development in the bone marrow of mice across the spectrum of physiological aging (young, aged, geriatric) compared with ELOVL2 MUT aged mice. Viably cryopreserved bone marrow samples from WT young, aged (MUT and WT), and WT geriatric mice were thawed and analyzed by flow cytometry using antibodies against murine CD19, CD79b, CD11b, and CD138. Samples were collected across n=9 separate experimental cohorts and analyzed in three separate flow cytometry batches. Results were combined by including individual replicate control samples across separate flow cytometry assays and data were normalized to the control values for each assay. A) Flow cytometry-based analysis of lymphocyte markers of the B cell lineage (CD19, CD79b) and myeloid markers (CD11b) were used to compare bone marrows from young-adult (3 months old, (n=5 female, n=7 male)), WT aged (18-20 months old, (n=3 female, n=15 male)), MUT aged (18-20 months old, (n=8 female, n=10 male)), and WT geriatric (28 months old, (n=4 male)) mice and frequency of live positive cells of each population are shown. For all panels, significance was determined using one-way ANOVA with parametric or non-parametric tests based on normality tests for each dataset (*p<0.05, **p<0.01, ***p<0.005, ****p<0.001; CD79b, CD11b – Kruskal-Wallis with Dunn’s multiple comparison post hoc test; CD19 – One-Way ANOVA with Tukey’s multiple comparison post hoc test). B) Representative flow cytometry gating strategy of CD19^+^/CD79b^+^ cells (live, single cells are shown). C) CD19^+^/CD79b^-/+^ and CD138^+^ markers were used to calculate the % frequency of progenitor-like and plasma cell populations in bone marrow samples. For all panels, significance was determined using one-way ANOVA with parametric or non-parametric tests based on normality tests for each dataset (*p<0.05, **p<0.01, ***p<0.005, ****p<0.001; CD19^+^/CD79b^-^, CD19^+^/CD79b^+^, CD138^+^: Kruskal-Wallis with Dunn’s multiple comparison post hoc test). D) Schematic diagram showing typical surface markers emerging during B cell progenitor maturation during the antigen-independent phase of B cell specification in the bone marrow (human cell markers shown based on^55–57^).

Further subpopulation analyses of the lymphoid lineage cells revealed that a mature CD19^+^CD79b^-^ population progressively decreased in frequency with early aging and to a greater extent with in WT geriatric mice, and that this advanced age feature was phenocopied in the *Elovl2*-MUT mouse bone marrows (Figure 2B). Surprisingly, a more primitive lymphoid progenitor-like population co-expressing CD79b was exclusively depleted in *Elovl2-*MUT and WT geriatric bone marrow versus aged WT controls (Figure 2B, C), as predicted by our bulk RNA-seq analyses. In contrast, plasma cells overall were much rarer (<2% of live cells in the BM) and were only found to trend downwards in the WT geriatric mice compared to other groups. Together, these results suggest that age-related and ELOVL2-associated lymphoid deficits may originate in a more primitive B-cell progenitor population that primarily impacts B cell development (Figure 2D) compared with other progeny of the lymphoid lineage.

Since organismal aging is known to frequently exhibit sex-specific differences and most females have longer average lifespans than males, we also explored the extent to which sex might influence the observed differences. Stratification of the flow cytometry data by sex revealed that there were no significant differences between sexes at each age or genotype studied, but that the overall trends remained the same for each sex when considered as separate cohorts, with a slightly stronger mutation-related effect observed in male mice (Figure S1C), although more female mice will need to be evaluated to confirm this observation.

Together, these data suggest that loss of a CD19^+^CD79b^+^ progenitor-like population might be a unique feature of advanced age that is recapitulated by systemic alterations in lipid metabolism in the setting of *Elovl2* deficiency.

### The aging bone marrow lipidome displays widespread biophysical and metabolic changes that are accelerated in the setting of ELOVL2 deficiency

Lipid biosynthesis of omega-3 (n-3) and omega-6 (n-6) PUFAs is a multi-step process involving specific desaturase, elongase, and β-oxidation enzymes (Figure 3A). This sequential process generates the myriad FA species necessary for the production of more complex lipids used in the generation of cellular membranes (predominantly phosphatidylcholine, PC) and energy storage (predominantly triglycerides, TG), among other cellular components.^29^ ELOVL2 is solely responsible for an essential elongation step that facilitates the endogenous production of DHA (Figure 3A). The C234W mutation in the ELOVL2 MUT mice selectively inactivates enzymatic activity of ELOVL2 required to process 22-carbon PUFAs. This results in a depletion of C24:5 and C22:6 (also known as DHA) species, while retaining elongase activity for other substrates common for ELOVL2 and the related enzyme ELOVL5.^7, 11^

**Figure 3:**
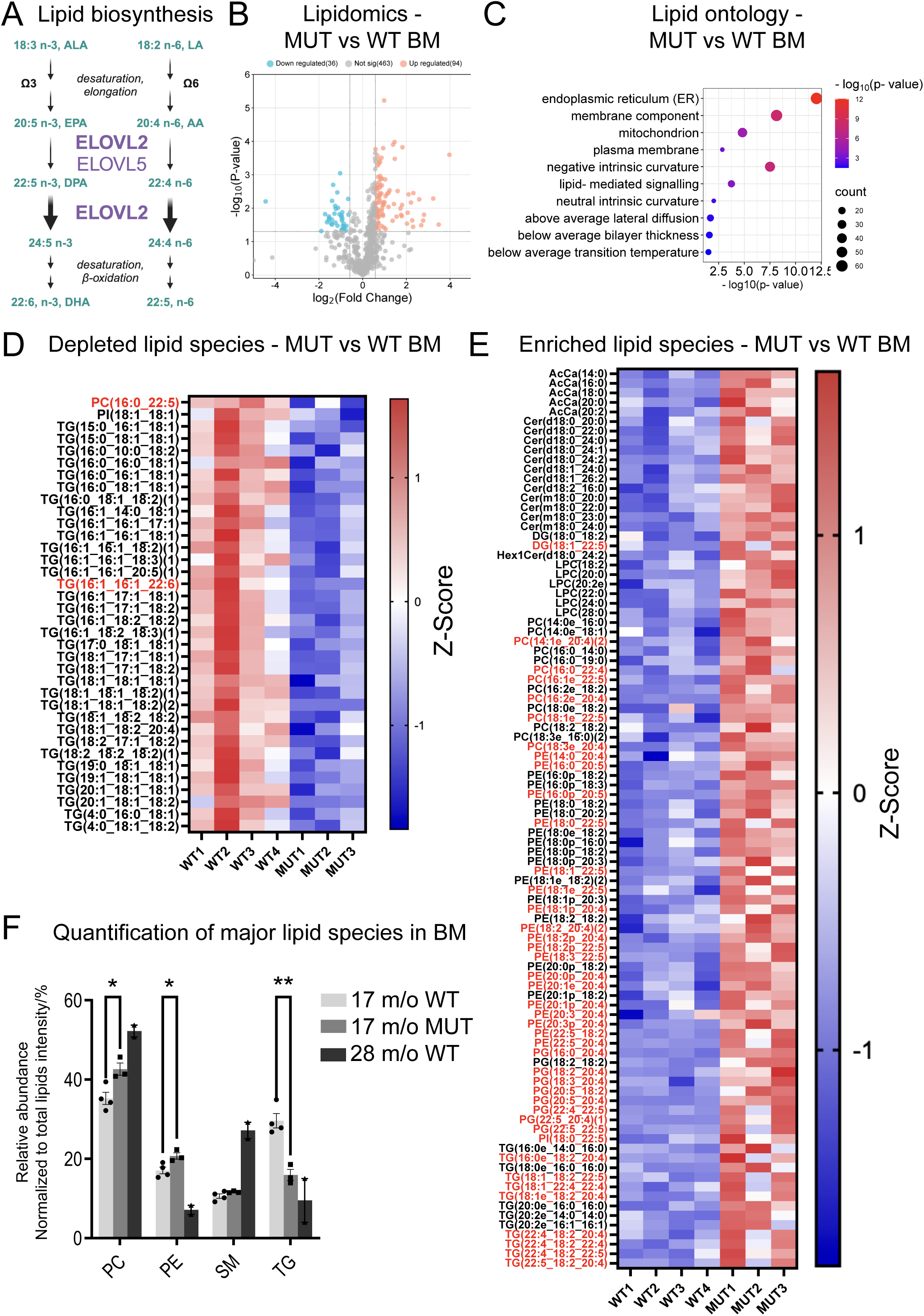
Global lipidomic analyses of total lipid class profiles and granular lipid species analyses in the bone marrow of MUT aged versus WT mice. Total bone marrow cells from aged (17 months old) WT (n=4 male) and MUT (n=3 male) mice were subjected to lipidomic analyses. Differentially expressed lipid species were analyzed by volcano plot, lipid ontology, and quantitation of individual lipid and total lipid class relative abundance. A) Schematic diagram of lipid biosynthesis pathway showing synthesis of omega-3 (n-3) and omega-6 (n-6) PUFAs, along with the specific elongation steps that ELOVL2 mediates and its substrate fatty acids (C20:5 n-3, C20:4 n-6) and its direct further downstream fatty acid products (C22:5 n-3, C22:4 n-6, C24:5 n-3, C24:4 n-6, C22:6 n-3, 22:5 n-6). B) Volcano plot visualizing significantly changed lipid species in aged MUT versus WT bone marrow samples (36 downregulated, 94 upregulated). C) Lipid ontology analyses comparing aged MUT versus WT bone marrow samples (D-E) Heatmaps showing differential lipid abundance (fold change >1.5, p<0.05) including significantly depleted lipid species (D) showing loss of ELOVL2 products (denoted in red) and significantly enriched lipid species (E) showing accumulation of ELOVL2 substrates (denoted in red). F) Relative abundance of major lipid species in aged MUT versus WT mouse bone marrow samples. Two additional samples from WT geriatric mice (28 months old) are shown for comparison. *p<0.05 and **p<0.01 compared to samples from WT aged (17 m/o) mice.

To generate a detailed lipid profile of accelerated bone marrow aging in ELOVL2-MUT mice, global lipidomics analyses were performed on 5×10^6^ total bone marrow cells per mouse from WT and MUT aged (17 months old) and WT geriatric (28 months old) animals. Volcano plot visualization and lipid ontogeny analyses of lipid species with differential abundances revealed that lipids involved in endoplasmic reticulum (ER) function, membrane and plasma membrane components, and lipid-mediated signaling, were the most significantly altered in MUT aged versus WT aged bone marrows (Figure 3B, C). Notably, a more granular analysis of all changed lipids revealed upregulation of diverse C22:5-containing lipid species in MUT aged mouse bone marrow, including PC (membrane-forming) species (Figure 3D, E). This is consistent with an accumulation of lipids containing C22:5 n-3 (or C20:5 n-3) molecules, which are the specific substrate that is unable to be processed by enzyme-deficient MUT ELOVL2. In contrast, the majority of downregulated lipid species were TG molecules (energy-storing), one of which contains a C22:6 lipid which is a likely product of normal ELOVL2 enzymatic function occurring only in WT mice. In addition, one C22:5 lipid species was identified as upregulated, which could potentially represent a product of ELOVL2 processing in the n-6 pathway, as our lipidomic technique cannot accurately distinguish between specific n-3 and n-6 species. Combined analyses of all major and minor lipid species demonstrate a global upregulation of PC species and loss of TG species in MUT versus WT aged mice (Figure 3F). Analysis of a few available WT geriatric mouse samples showed similar alterations in levels of PC and TG species, which resembled much younger MUT mice (Figure 3F). Minor lipid species were mostly unchanged across the groups, with the exception of predominantly saturated FA-containing acetyl-carnitines (AcCa) and some ceramide species, which are used by mitochondria as energy, and lipid rafts in the plasma membrane, respectively (Figure S2A).^29^

To extend these analyses and explore whether global lipid changes in the bone marrows of MUT versus WT aged mice might resemble the physiological trajectory of bone marrow aging at advanced ages, WT geriatric versus WT aged bone marrows were also compared to each other (Figure S2B, C). Lipid ontology analyses of WT bone marrow cells revealed that physiological aging in geriatric mice impacted similar pathways (ER, plasma membrane, lipid-mediated signaling) as those that were changed in MUT mice at younger ages (Figure S2C), further supporting the possibility that ELOVL2-MUT mice recapitulate diverse molecular, cellular, and biophysical features of organismal aging, which occur in this model at an accelerated rate.

To confirm that there is no other endogenous source of DHA or other products of ELOVL2 PUFA elongation in the circulation of mutant animals, levels of total FAs, free FAs, along with lipid species alterations were assessed in MUT versus WT aged mouse plasma samples (Figure S2D, E). Among 538 total analytes, 25 lipid species were upregulated and 37 lipid species were downregulated in MUT aged mice compared with WT age-matched controls. A striking loss of all n-3 22-carbon and longer products of ELOVL2 activity was observed in total FA analysis of MUT versus WT aged mouse plasma, coupled with an accumulation of upstream substrates in the n-3 pathway (e.g., 22:5-1) (Figure S2D). Free FAs represent a much smaller proportion of lipids in plasma, however the same types of changes were observed (Figure S2D). As expected, and similar to what has been previously reported in *Elovl2* knockout mouse plasma,^30^ lipid species containing C22:5 (or C20:5) components were predominantly enriched, while lipid species containing C22:6 components were predominantly depleted in MUT versus WT aged mouse plasma (Figure S2E).

Together, total lipid class profiles and granular lipid species analyses showed down-regulation of select PUFAs and accumulation of shorter-carbon FAs and saturated FAs in the bone marrow of MUT aged mice, supporting the possibility that *Elovl2* mutation may mimic the age-related disruption of biosynthesis of unsaturated FAs and disrupt metabolic function during blood cell development.

### ELOVL2 is expressed in a rare subset of CD34^+^ human hematopoietic stem and progenitor cells and is depleted in bone marrow from elderly individuals

To further explore the clinical relevance of these findings and elucidate connections between lipid homeostasis and lymphoid specification in human samples, we analyzed public datasets along with a unique human cell dataset^18^ to characterize *ELOVL2* expression and its correlation with lymphoid cell markers at single-cell resolution in CD34^+^ human HSPCs spanning gestation, maturation, and aging. An overall loss of lymphoid-primed progenitors with age was observed in this prior study, and at the gene expression level, these age-related changes appeared to be ingrained within the HSC compartment.^18^ Consistent with a depletion of lymphoid-primed progenitors in elderly individuals, gene-specific profiling of *CD79B* in sorted CD34^+^ cells revealed a stable population of *CD79B*-positive cells across development from childhood and adolescence, through adulthood (17-53 years old, average age 34.5 years, n=6), with a dramatic loss observed in bone marrow samples from elderly individuals (62-77 years old, average age 71.7 years, n=3) (Figure 4A). This corresponded with a decrease in lymphoid-primed HSPC from 20.4% in adults to 8.8% in elderly individuals (Figure 4B), validating the results from the mouse model showing that CD79B could represent a unique indicator of lymphoid immune cell fitness during aging. Notably, analysis of *ELOVL2* in these different progenitor compartments revealed the presence of a subset comprising 6.7% of all CD34^+^ HSPCs that express *ELOVL2* in healthy adult bone marrow (Figure 4C). Most of these cells were within sub-compartments denoted as uncommitted, HSC, or lymphoid-primed, and they were almost undetectable in CD34^+^ cells isolated from elderly bone marrow samples (0.8% of total CD34^+^ cells, Figure 4D).

**Figure 4:**
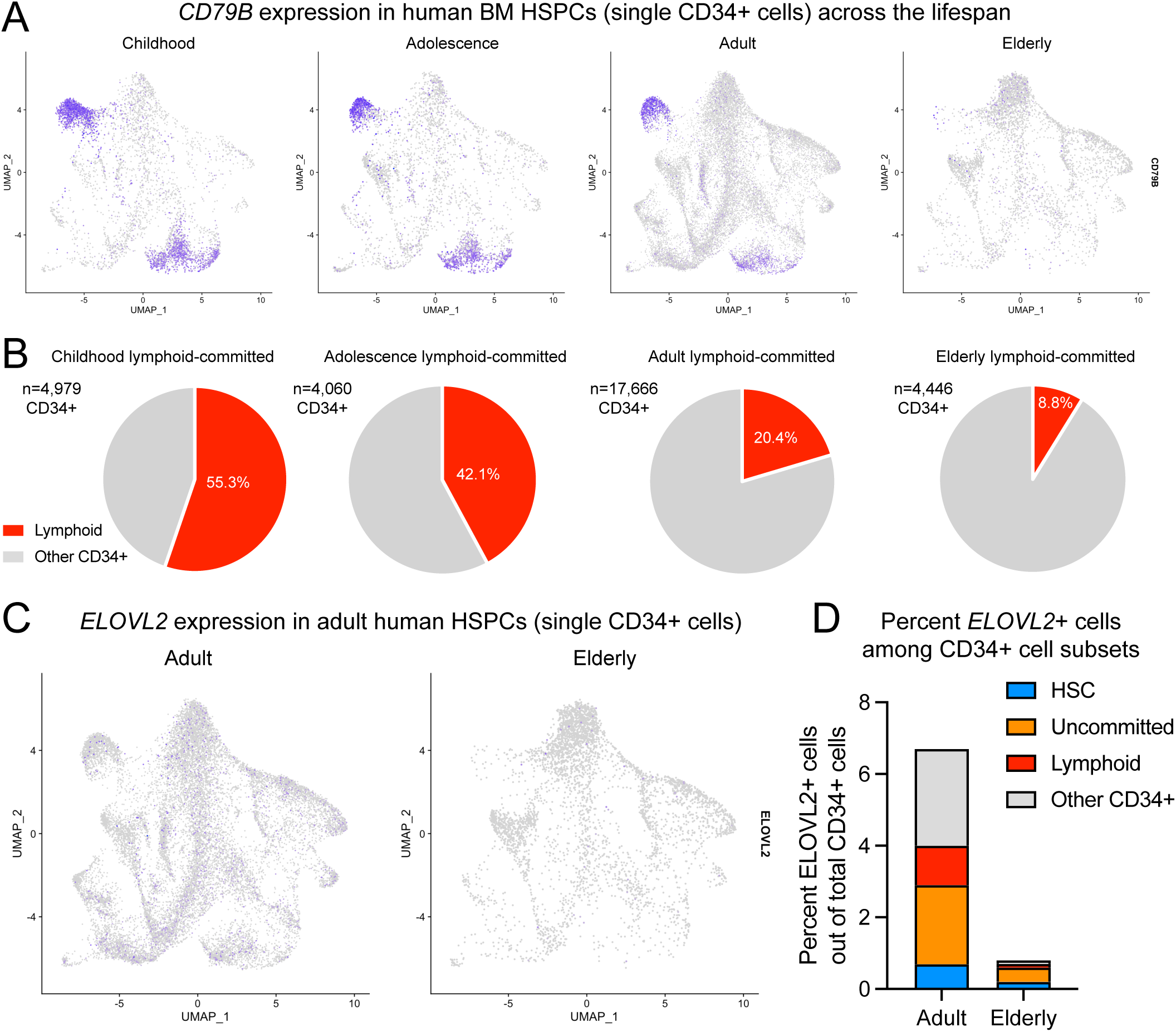
Single-cell analysis of lymphoid-committed cells and ELOVL2^+^ subsets of human CD34^+^ human hematopoietic stem and progenitor cells across the spectrum of human development and aging. Previously described datasets from single-cell RNA-sequencing studies of human CD34+ cells were subjected to gene-specific analysis of lymphoid marker genes and ELOVL2.^18, 19^ A) UMAP plots showing relative CD79B expression in single sorted CD34^+^ cells throughout development from childhood (2-4 years old) to adolescence (10-12 years old), adulthood (ages 17-53), and elderly individuals (ages 62-77). B) Quantification of lymphoid-committed hematopoietic stem and progenitor cell frequencies (based on gene programs described in^18^). C) UMAP plots showing relative ELOVL2 expression in single sorted CD34^+^ cells in adult (n=6) versus elderly (n=3) human subjects as in panel (A). D) Quantification of changes in frequency of ELOVL2^+^ cells among CD34^+^ cell subsets in elderly versus adult human subjects.

## Discussion

A bias toward myeloid cell development is one of the most well-characterized phenotypes of the aging immune system and is frequently linked to a pre-disposition to developing CHIP or clonal cytopenias of undetermined significance (CCUS) leading to pre-malignant or malignant myeloid disorders. However, mammalian aging is typified by a complementary loss of lymphoid populations^18, 19, 31^ and aging often occurs in the setting of diverse age-related metabolic disorders, which have been associated with defective lymphopoiesis. For example, obesity has been linked to impaired lymphoid cell development, and a high-fat diet results in rapid changes to B cell development,^32, 33^ however the precise FA metabolism pathways that impair immune system function during aging have remained unclear.^34^ In the present study, we uncovered a rare subpopulation (<7%) of human bone marrow CD34^+^ HSPCs that express an essential enzymatic regulator of LC-PUFA synthesis, ELOVL2. This population is almost completely lost in bone marrow samples from elderly individuals when compared to healthy adult donors, which are also severely depleted of lymphoid-primed subpopulations.^18^

While most previous aging bone marrow studies have focuses on altered lipid accumulation and metabolism in non-hematopoietic cells of the bone marrow niche,^35, 36^ some more detailed studies of the metabolome^37^ and transcriptome of aged versus young mice have reported alterations in fatty acid metabolism and disruption of B cell development and stem cell related pathways (e.g., Wnt/b-catenin and JAK/STAT signaling).^38^ In this context, we sought to explore whether ELOVL2 enzymatic deficiency might be associated with features of physiological bone marrow aging including altered HSPC development. In genetically engineered mouse model studies, we demonstrated that systemic enzymatic deficiency of ELOVL2 is associated with dramatic loss of B cell populations expressing markers of the pro- and pre-B cell stages of development. First, whole transcriptome RNA-sequencing based analyses with qRT-PCR validation identified loss of *Cd79b*, *Cd19* gene expression along with downregulation of key lymphopoiesis regulatory transcription factors *Pou2af1*, *Pax5*, *Spib* and *Irf4* as features of an accelerated aging phenotype in the marrow of ELOVL2-deficient mice. Flow cytometry analyses in expanded cohorts of mice confirmed these observations and further revealed that a specific subpopulation of CD79B/CD19 double-positive cells was selectively depleted in aged MUT mouse BM compared with WT controls, and that these changes phenocopied the molecular and cellular alterations observed in geriatric WT mouse BM. Together, these results suggest that impaired synthesis of ELOVL2-dependent LC-PUFA species could drive an aging-associated maturation defect in an early B cell progenitor compartment.

Proper control of FA metabolism is essential for stem cell self-renewal^39, 40^ in part because of their high bio-energetic needs and demand for the raw materials (FAs) necessary for plasma membrane and organelle production.^39, 41^ ELOVL2 is the rate-limiting enzyme responsible for endogenous production of the LC-PUFA DHA (22:6, n-3). Previous studies in an *Elovl2* knockout mouse model demonstrated that DHA is endogenously produced and is essential for lipid homeostasis throughout the organism.^30^ In the present study, we observed a similar systemic loss of DHA levels in ELOVL2-mut mice as was previously shown in *Elovl2-*deficient mice, indicating that the ELOVL2 mutation accurately phenocopies overt genetic loss of this enzyme. The present study reveals that the lipidome of ELOVL2 MUT bone marrow samples was dramatically remodeled compared to WT controls, with loss of products of ELOVL2 metabolism in BM and plasma samples, and aged ELOVL2 MUT mouse BM at 18-20 months old showed similar lipidomic profiles to geriatric mouse samples (27 months and older). The specific lipid species changed in the MUT and geriatric WT mouse BM suggests biophysical changes to the plasma membrane and other membrane-containing cellular components, along with energy storage. Importantly, loss of ELOVL2-dependent LC-PUFA, or alteration in their ratio relative to the abundance of shorter or saturated FA species, has the potential to completely disrupt the plasma membrane structure and fluidity characterized by a stiffer membrane composition.

The plasma membrane is where key components of the pre-B cell receptor (pre-BCR) and BCR are assembled, which are essential for B cell maturation. During the antigen-independent early phase of B cell development from lymphoid-primed hematopoietic progenitors (CD34^+^CD10^+^),^42^ which occurs in the BM, expression and heterodimerization of CD79B (Igβ) along with CD79A (Igβ) are among the key initiating steps of pre-B cell receptor (BCR) complex generation. CD79A and CD79B are vital in the transition of pro-B cells to pre-B cells^43^ and selective expansion of pre-B cells,^44, 45^ among other essential roles they play in B cell maturation.^46^ Disruption of CD79A or B completely blocks B cell development in mice,^47, 48^ and CD79B, but not CD79A has been reported to have the ability to form a homodimer that is transported to the developing B-cell surface without any other BCR components present.^49^ Furthermore, during B cell development, physical translocation of the B cell receptor (BCR) complex from the lipid membrane to lipid rafts is required for receptor cross-linking, antigen-mediated signaling, and proper B cell maturation.^50^ Although a limitation of the present study is that we are unable to evaluate the direct mechanism through which ELOVL2 blocks B cell development, based on the molecular, cellular, and lipidomic alterations we describe in the present study, we predict that ELOVL2-deficiency and advanced age may promote structural plasma membrane alterations that alter the activity of vital transmembrane proteins. This, in turn could impinge upon pre-BCR and BCR assembly and signaling and act as a biophysical block to lymphopoiesis in physiologic or accelerated aging, which will be explored in future studies.

Although the BM aging phenotypes reported here involve loss of B-cell populations expressing CD79B, the findings have important implications for conditions that involve aberrant activation of these pathways, especially considering that metabolic gatekeeper functions during B-cell development have been implicated in safeguarding against both autoimmune diseases and B-cell malignancies.^51, 52^ For example, CD79B appears to harbor additional signaling-related functions which drive the pro- to pre-B cell transition through activation of Bruton’s tyrosine kinase (BTK).^46^ Notably, activation of BTK signaling is an essential feature of some B-cell malignancies such as chronic lymphoid leukemia (CLL) and Waldenstrom’s macroglobulinemia (WM), and inhibitors of BTK (i.e., ibrutinib) are standard of care therapies for these cancers. In addition, CD79B transcripts are significantly more abundant than CD79A (4-fold increase) in some lymphoid cancer cell lines, suggesting that CD79B could act as a putative tumor promoter in leukemic B-cells.^53^ Since loss of ELOVL2 activity regulates the abundance of CD79B- expressing cells in mice, future studies could also focus on exploring ELOVL2-dependent lipid synthesis pathways and/or PUFA-targeted dietary interventions as novel therapeutic targets for the prevention or treatment of B-cell malignancies.

Taken together, we have uncovered the novel mechanism of age-related HSPC decline driven by disrupted lipid homeostasis and have identified novel cellular indicators of accelerated immune aging (CD79B). These studies provide new insights as well as molecular targets which could have future applications for therapeutic intervention in clinical conditions related to aging and cancer.

## METHODS

### Animal Husbandry

Mice were housed under a 12-hour light/dark cycle at 22±2°C with free access to standard laboratory diet chow (Teklad 2020x) and water. All animal procedures were performed in accordance with the UC-Irvine Animal Care and Use Committee guidelines and IACUC protocols (#AUP-23-064). Male and female mice (C57BL/6), young (2-3 months old), aged (18-22 months old), and geriatric (27-28 months old) were used for experiments.

### Murine Bone Marrow Tissue Harvesting and Processing

Bone marrow was harvested from the femurs and tibias of euthanized mice by flushing the marrow cavity with Staining Media (STM: HBSS 1x, 2% FBS-heat inactivated (HI), 2mM EDTA) using a 25-gauge needle and 10ml syringe. The harvested bone marrow cells were then passed through a 70-μm cell strainer to remove bone fragments and aggregates. Cells were counted and centrifuged at 300g for 5 minutes at 4°C, and the pellet was divided among the different fractions for subsequent analyses and processing. The RNA fraction was resuspended in RLT buffer with 2-Mercaptoethanol (BME >99%) and stored at –80°C. Each sample’s lipidomic pellet was flash-frozen, containing 5 million cells to account for harvesting day biases and ensuring a consistent amount of lipids. The flow cytometry fraction was stored for cryopreservation in liquid nitrogen in DMSO/FBS-HI (10%/90%). All fractions were stored until further processing. In some experiments, a greater number of male MUT mice were used for analyses because female animals of the MUT line were reserved for breeding, however whenever possible we also compared results stratified by sex to assess for any sex-specific differences.

### Bulk RNA-seq

Total RNA was extracted from bone marrow cells using the AllPrep DNA/RNA Mini Kit (Qiagen, #80204) and RNeasy Mini Spin Columns (Qiagen, #1112543) according to the manufacturer’s instructions. RNA integrity, library preparation and sequencing were carried out by the UCSD IGM Genomics Center. Total RNA was assessed for quality using an Agilent Tapestation 4200, and samples with an RNA Integrity Number (RIN) greater than 8.0 were used to generate RNA sequencing libraries using the Illumina® Stranded mRNA Prep (Illumina, San Diego, CA). Samples were processed following manufacturer’s instructions. Resulting libraries were multiplexed and sequenced with 100 basepair (bp) Paired End reads (PE100) to a depth of approximately 25 million reads per sample on an Illumina NovaSeq 6000. Samples were demultiplexed using bcl2fastq Conversion Software (Illumina, San Diego, CA).

### Gene Set Enrichment Analysis

Raw data were processed and analyzed using a standardized pipeline including quality control, alignment to the mouse genome (mm10), and differential expression analysis with Rosalind Platform (https://www.rosalind.bio/). Gene-set enrichment analysis (GSEA) was performed to identify significantly enriched biological pathways. Pre-ranked gene lists based on log2 fold changes from the RNA-seq data were analyzed against KEGG pathways. False discovery rates (FDR) were calculated to identify statistically significant pathways (FDR < 0.05).

### Bone Marrow Aging-Associated Molecular Signatures

Differential expression analysis from bulk-RNA-seq data was used to identify ELOVL2-mutation and aging-regulated genes in bone marrow samples. Mutant ELOVL2-associated molecular signatures were defined as genes showing significant expression changes between aged MUT and aged WT mice. These signatures were further analyzed for pathway enrichment and cross-referenced with literature data to validate bone marrow/immune system aging relevance.

### cDNA Preparation and Quantitative RT-PCR

RNA samples from mouse bone marrows were purified as described above for bulk RNA-sequencing and eluted by adding 30 µL of sterile, RNase-free water directly onto the membrane, incubating for 3–5 minutes, and centrifuging at 14,000 rpm for 1 minute. RNA was stored at −80°C. For cDNA synthesis, RNA samples were thawed on ice, and concentrations were measured using a NanoDrop spectrophotometer (ThermoFisher). Calculations were performed to determine RNA input for cDNA synthesis, targeting consistent output despite varying RNA inputs. The reaction mix was assembled by sequentially adding water, RNA, and VILO mastermix (MM), followed by brief centrifugation. The cDNA synthesis was conducted using the BioRad iCycler with the VILO protocol (20 µL reaction, 1 cycle, 4 steps) without a hot start. Reactions ran for 30 minutes, and cDNA was stored at −20°C. Internal positive control (human cell line reference) was included for plate normalization. Negative control (no template control) consisted of 16 µL of water and 4 µL of MM without RNA.

### Quantitative Real-Time PCR (qRT-PCR)

qRT-PCR was performed using TaqMan assays with primers targeting murine *Elovl2, Cd79b,* or *Pou2af1* (FAM) and *Hprt* (VIC) as a housekeeping gene (ThermoFisher Taqman assay IDs: *Elovl2* Mm00517086_m1*, Cd79b* Mm00434143_m1, or *Pou2af1* Mm00448326_m1*, Hprt* Mm03024075_m1). Assays were run in technical duplicates with each gene multiplexed with the housekeeping gene in individual wells. cDNA samples and primers were retrieved from −20°C storage, and master mix was kept on ice during preparation. To prepare the qPCR mix, ultra-pure distilled water was mixed with the primers, and the TaqMan enzyme master mix. The prepared mix was dispensed into a 96-well Applied Biosystems plate, pre-cooled on a cold plate holder, followed by the addition of 1 µL of each cDNA sample. After covering with film, the plate was centrifuged at 1300 rpm for 3 minutes at 4°C. The qPCR plate was analyzed using a QuantStudio3 instrument (ThermoFisher), and data were analyzed in Excel and graphs plotted using GraphPad Prism.

### Flow Cytometry and Immunophenotyping Panels

#### Sample Preparation

Bone marrow cells from *Elovl2* mutant and wildtype mice were thawed and resuspended in Staining Media (STM). Cells were counted and adjusted to a concentration of 2 million cells per sample. For viability assessment and subsequent antibody staining, two different panels (Plate A and Plate B) were prepared as per the immunophenotyping protocols provided.

#### Staining Protocol

##### Plate A (Lymphoid/Myeloid Panel without Fixation)

Cells were washed with PBS at room temperature and centrifuged at 300g. Viability staining was performed by incubating the cells in Viobility Fixable Dye (405/520) (cat# 130-130-421 Miltenyi) at room temperature, in the dark, for 15 minutes (1 µL of stain to 100 µL PBS per well). Cells were washed twice with ice-cold STM (HBSS 1x, 2% FBS-HI, 2mM EDTA) and centrifuged at 300g. Antibody staining was conducted by adding fluorochrome-conjugated antibodies against CD11b (viobright FITC, (Miltenyi, # 130-113-805)), CD19 (PE, (Miltenyi, # 130-112-035)), and CD79b (APC, (Miltenyi, # 130-119-426)) diluted 1:100 in STM. The cells were incubated for 10 minutes in the dark at 4°C, followed by two washes with ice-cold STM.

##### Plate B (Plasma Cell Panel with Fixation)

Cells underwent viability stain and cell-surface staining against CD19 (viobright 515, (Miltenyi, # 130-112-040)), CD138 (PE, (Miltenyi, # 130-120-810)), as described in Plate A. Following that, cells were fixed with True-Nuclear™ 1X Fix Concentrate (BioLegend, # 73158) for 60 minutes at room temperature in the dark. The cells were washed and permeabilized using True-Nuclear™ 1X Perm Buffer (BioLegend, # 73162). Intracellular staining was performed using fluorochrome-conjugated antibodies against IRF4 (AF647, (BioLegend, 646408)) in 1X Perm Buffer for 30min in the dark, followed by three washes with 1X Perm Buffer.

#### Flow Cytometry

The cells were resuspended in cell staining buffer and acquired on a BD FACSymphony™ A1 Cell Analyzer supported by BD FACS Diva Software.

#### Data Acquisition and Analysis

For each sample, a minimum of 30,000 events were recorded. Data acquisition settings and gating strategy (BM cells+, Single Cells+, Live Cells+, FMOs for the antibody staining) were consistent among experiments to ensure comparability. Data analysis was conducted using FlowJo, with gates set to exclude doublets and debris, followed by gating on live cells based on viability dye exclusion. Subsequent gates were applied to identify lymphoid, myeloid, and plasma cell populations based on specific marker expression.

### Lipidomics

Lipid fractions were extracted from bone marrow samples using a modified Bligh and Dyer method, followed by hydrolysis for total acids analysis from bone marrows of WT and MUT female and male mice, as previously described.^54^ To ensure consistency of results, for each sample, 5×10^6^ total bone marrow cells were used for lipidomic analyses, and values were normalized to total lipid intensity.

### Lipidomic analysis

#### Lipid extraction

Lipid extractions were performed according to the methodology of Bligh and Dyer^18^. In brief, the tissue was homogenized in 200 μL water, transferred to a glass vial, and 750 μL 1:2 (v/v) CHCl_3_: MeOH was added and vortexed. Then 250 μL CHCl_3_ was added and vortexed. Finally, 250 μL ddH_2_O was added and vortexed. The samples were centrifuged at 3000 RPM for 5 min at 4 °C. The lower phase was transferred to a new glass vial and dried under nitrogen stored at −20 °C until subsequent lipid analysis.

#### LC-MS/MS

Separation of lipids was performed on an Accucore C30 column (2.6 μm, 2.1 mm × 150 mm, Thermo Scientific). The Q Exactive MS was operated in a full MS scan mode (resolution 70,000 at m/z 200) followed by ddMS2 (17,500 resolution) in both positive and negative mode. The AGC target value was set at 1E6 and 1E5 for the MS and MS/MS scans, respectively. The maximum injection time was 200 ms for MS and 50 ms for MS/MS. HCD was performed with a stepped collision energy of 30 ± 10% for negative and 25%, 30% for positive ion mode with an isolation window of 1.5 Da.

#### Data analysis and post-processing

Data were analyzed with LipidSearch 4.2.21 software. Only peaks with molecular identification grade: A or B were accepted (A: lipid class and fatty acid completely identified or B: lipid class and some fatty acid identified).

### FA analysis

#### FA extraction

1 μg 6Z,9Z,12Z,15Z,18Z-heneicosapentaenoic acid (FA 21:5, Cayman, USA) was added as internal standard. Lipids were extracted by the Bligh-Dyer method followed by hydrolysis and extraction of total fatty acids. Briefly, 750 μL ice-cold 1:2 (v/v) CHCl_3_:MeOH was added directly to the sample and the sample was vortexed. Then, 250 μL CHCl_3_ was added and mixed well. 250 μL ddH_2_O was added and vortexed well to produce a two-phase system. After centrifuging at 3000 RPM for 3 min, the bottom phase was collected and evaporated under a constant stream of nitrogen. To recover the total fatty acids, 720 μL acetonitrile and 10 μL hydrochloric acid were added to the dried film and the sample was kept at 95 °C for 1 h. Then, 1 ml of hexane was added to the sample and vortexed. The upper phase containing total fatty acids was transferred to a new tube and evaporated under nitrogen. The total fatty acids were dissolved in acetonitrile-isopropanol (50:50, v/v) for further LC-MS Analysis.

#### LC-MS/MS

Separation of VLC-PUFAs was achieved on an Acquity UPLC® BEH C18 column (1.7 μm, 2.1 mm × 100 mm, Waters Corporation). The Q Exactive MS was operated in a full MS scan mode (resolution 70,000 at m/z 200) in negative mode. For the compounds of interest, a scan range of m/z 250–800 was chosen. The identification of fatty acids was based on retention time and formula.

### Single-cell Analysis of Lymphoid Markers and *ELOVL2* Expression in Primary Human HSPCs

Relative lymphoid cell frequencies and counts of cells expressing *CD79B* and *ELOVL2* were extracted from a previously described single-cell RNA-seq dataset of human CD34^+^ HSPCs.^18, 19^ *CD79B* and *ELOVL2* expression was overlayed on UMAPs made from CD34^+^ bone marrow cells at various points along the human lifespan.

### Statistical Analyses

Data were analyzed using GraphPad Prism (Prism 10, GraphPad Software). Normality of the data was assessed using the Shapiro-Wilk test. Depending on the distribution of the data, parametric or non-parametric statistical tests were applied accordingly. For normally distributed data, one-way ANOVA followed by Tukey’s multiple comparison test was used to compare differences between groups. For non-normally distributed data, the Kruskal-Wallis test followed by Dunn’s multiple comparison test was applied. A *p*-value of <0.05 was considered statistically significant.

## Data Availability

Mouse bone marrow whole transcriptome sequencing data files and raw lipidomics datasets will be made publicly available after formal publication. Human CD34^+^ single cell datasets are available in GEO under accession #GSE189161.^18, 19^

## Acknowledgements

S.V. is supported by a CIRM fellowship (EDUC4-12804) and L.A.C. is a Scholar of The Leukemia & Lymphoma Society and a Sanford Stem Cell Institute Stellar Faculty Award recipient. Research in the Crews Lab is supported by NIH/NCI R37CA252040 and in part by NIH/NCI P30CA023100, the International Myeloma Foundation, the International Waldenstrom’s Macroglobulinemia Foundation, the Leukemia Research Foundation, the UC San Diego Health Sciences Senate, and the UC San Diego Sanford Stem Cell Institute. Research in the DS-K laboratory is funded by NIH/NEI U01EY034594, and in part by NIH grant P30EY034070 and an unrestricted grant from Research to Prevent Blindness (New York, NY, United States) awarded to the Department of Ophthalmology, UC Irvine. H.L. is supported by NIH/NIDDK K08DK123414. The authors wish to thank Diane McCann, Anna Rapp, Rosslyn Farnan, Astha Patel, Elizabeth Diaz, Thon de Boer, Max Rempel, and Nga Phuong Nyugen for technical support, and Catriona H.M. Jamieson for helpful discussion. Bulk RNA-sequencing was conducted at the IGM Genomics Center, University of California, San Diego, La Jolla, CA, which is supported by NIH/NCI P30CA023100. This publication includes data generated at the IGM Genomics Center utilizing an Illumina NovaSeq X Plus that was purchased with funding from a National Institutes of Health SIG grant (#S10 OD026929). Flow cytometry data were collected at the UCSD Moores Cancer Center flow cytometry facility, which obtained a BD FACSymphony S6 through support from the NIH (S10OD032316). Some figures were generated using Biorender.

## Conflicts of interest

DS-K is a scientific advisor of Visgenx, Inc.

## Supplemental Information

### Supplemental Figure Legends

**Figure S1 (related to Main Figures 1 and 2):**
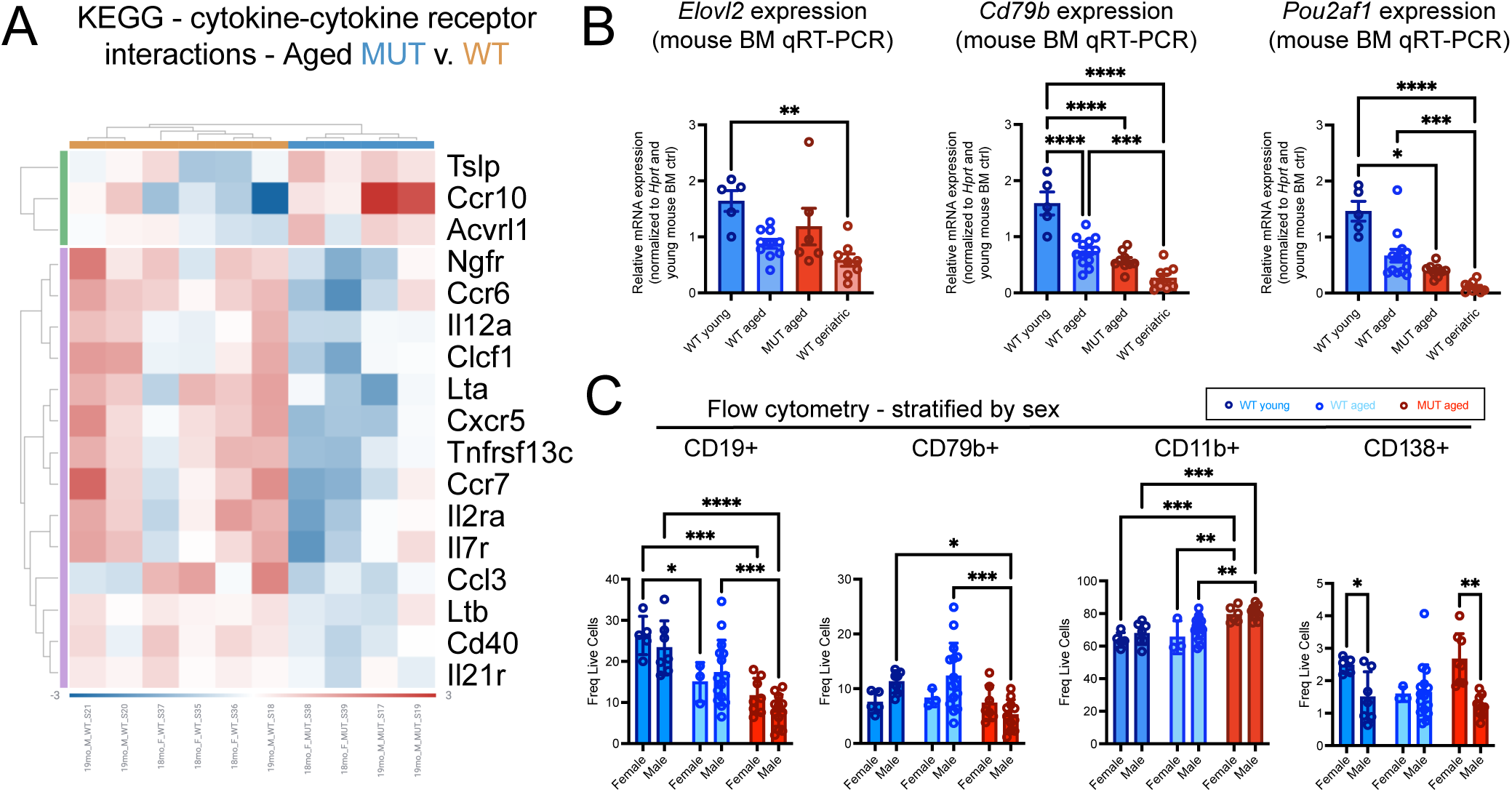
Validation of multi-omics analyses and sex-stratified protein expression changes in the bone marrow of WT young, aged, geriatric, and ELOVL2 MUT mice. Total bone marrow cells from 18-19 month old aged WT versus ELOVL2 C234W MUT mice were analyzed by whole transcriptome RNA-sequencing (as in Figure 1) with expanded analyses in additional samples for qRT-PCR and flow cytometry. A) Heatmap showing the relative expression levels of KEGG cytokine-cytokine receptor interactions pathway gene orthologs in aged MUT versus WT bone marrow samples. B) qRT-PCR analysis of *Elovl2, Cd79b,* and *Pou2af1* gene expression in an expanded cohort of mouse bone marrow samples including WT young, MUT aged, WT aged and WT geriatric. For all panels, significance was determined using one-way ANOVA with parametric or non-parametric tests based on normality tests for each dataset (*p<0.05, **p<0.01, ***p<0.005, ****p<0.001; *Elovl2*, *Pou2af1*: Kruskal–Wallis with Dunn’s multiple comparison post hoc test; *Cd79b*: One-Way ANOVA with Tukey’s multiple comparison post hoc test). C) Flow cytometry data from Main Figure 2 (A and C), stratified by sex. Since no young male samples were available for these analyses, the young condition was not plotted since n=5 were all females. For all panels, significance was determined using one-way ANOVA with parametric or non-parametric tests based on normality tests for each dataset (*p<0.05, **p<0.01, ***p<0.005, ****p<0.001; Cd19 and Cd79b: Two-Way ANOVA with Tukey’s multiple comparison post hoc test; Cd11b: Two-Way ANOVA with Šidak’s multiple comparison post hoc test; *p<0.05; Cd138: Mixed effect model with Dunn’s multiple comparison post hoc test; *p<0.05).

**Figure S2 (related to Main Figure 3):**
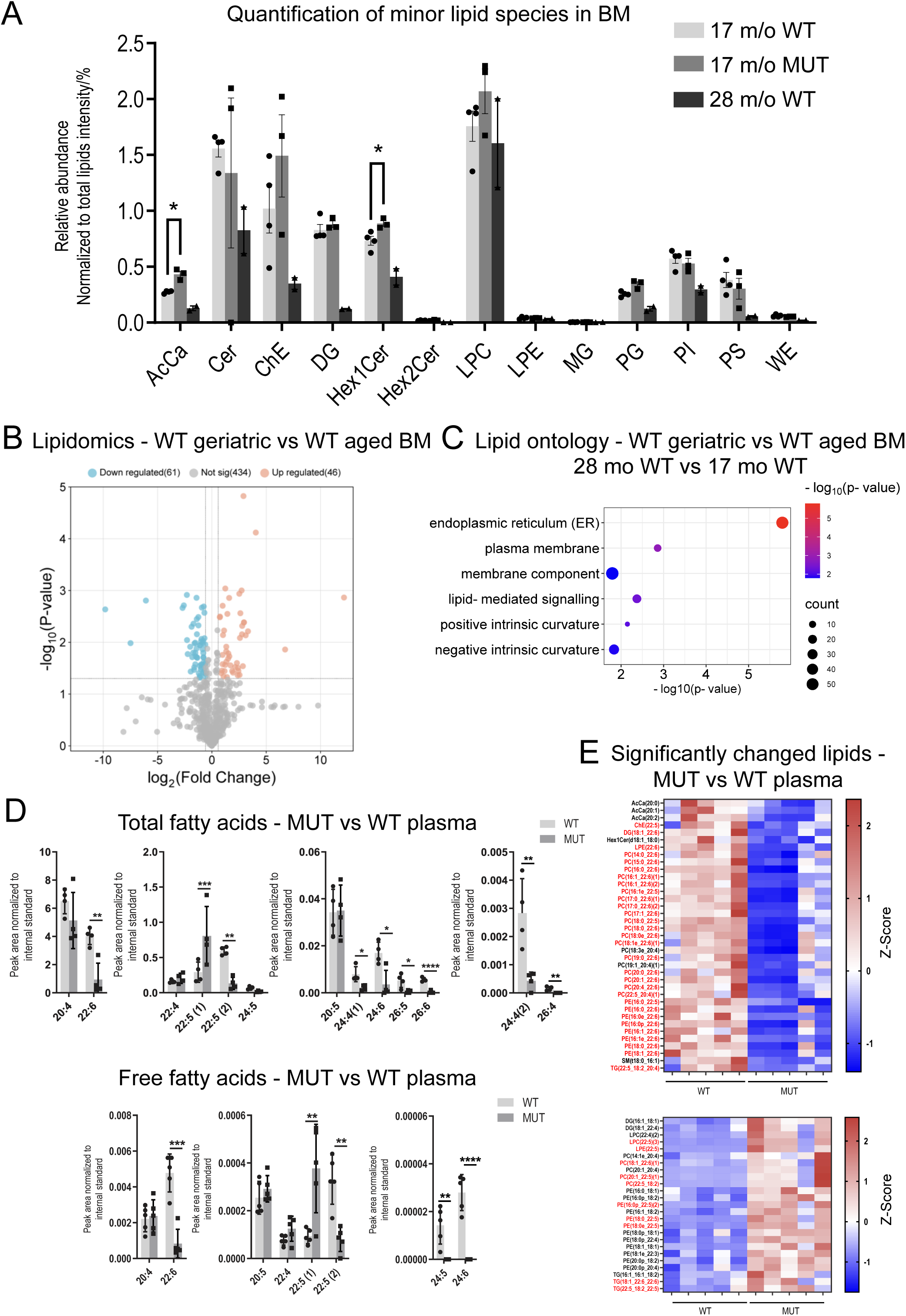
Global lipidomic analyses of the bone marrow of WT aged, WT geriatric, and MUT aged mice and fatty acid and lipidomic analyses of plasma samples from aged MUT versus WT mice. A) Relative abundance of minor lipid species compared among aged MUT vs. aged WT vs. geriatric WT. *p<0.05 compared to samples from WT aged (17 months old) mice. B) Volcano Plot indicates significant differential expression between geriatric WT vs. aged WT bone marrow samples. C) Lipid ontology analyses comparing WT geriatric versus aged bone marrow samples. D) Total and free fatty acids analyses in aged (18 months old) MUT vs WT plasma samples (*p<0.05 compared to WT aged plasma samples). E) Heatmaps showing differential abundance (fold change >1.5, p<0.05) of lipid species in aged MUT vs WT plasma (upper panel shows significantly downregulated lipids and lower panel shows significantly upregulated lipids in aged MUT plasma).

